# AimB is a small protein regulator of cell size and MreB assembly

**DOI:** 10.1101/655720

**Authors:** John N. Werner, Handuo Shi, Jen Hsin, Kerwyn Casey Huang, Zemer Gitai, Eric A. Klein

## Abstract

The MreB actin-like cytoskeleton assembles into dynamic polymers that coordinate cell shape in many bacteria. In contrast to most other cytoskeletons, few MreB interacting proteins have been well characterized. Here we identify a small protein from *Caulobacter crescentus*, AimB, as an <u>A</u>ssembly <u>I</u>nhibitor of <u>M</u>re<u>B</u>. AimB overexpression mimics inhibition of MreB polymerization, leading to increased cell width and MreB delocalization. Molecular dynamics simulations suggest that AimB binds MreB at its monomer-monomer protofilament interaction cleft. We validate this model through functional analysis of point mutants in both AimB and MreB, photo-crosslinking studies with site-specific unnatural amino acids, and species-specific activity of AimB. Together, our findings indicate that AimB promotes MreB dynamics by inhibiting monomer-monomer assembly interactions, representing a new mechanism for regulating actin-like polymers and the first identification of a non-toxin MreB assembly inhibitor.

## Introduction

Maintenance of proper cell size is an important physiological process for all organisms. Changes in cell size are often strongly coupled to cell fitness in laboratory evolution experiments^1^, and mutations that affect cell size can be highly adaptive^2^. Cell size is also dynamically regulated, as in the example of rod-shaped bacteria whose dimensions are altered by environmental factors such as nutrient availability^3^. A recent study developed a biophysical model in which cell size is determined by the relative rates of surface area and volume synthesis; upon nutrient upshift, the increased rate of cytoplasmic synthesis reduced the surface area-to-volume ratio via an increase in cell width^4^. However, the molecular regulators of cell size remain largely unclear in most bacterial species.

Bacterial cell shape determination requires enzymes that directly synthesize and crosslink peptidoglycan chains in the periplasm and cytoskeletal factors that localize the activity of these enzymes. The actin homolog MreB serves this cytoskeletal function for cell elongation. Studies from *Escherichia coli* and *Bacillus subtilis* show that MreB forms filaments that localize and move^5–7^ along the membrane based on the local cell geometry^8^ and recruit cell wall enzymes to insert new cell wall and change the shape of those sites, resulting in a feedback loop that establishes rod shape^9^. In this model, MreB must dynamically assemble and disassemble to sample multiple cellular regions over time, and indeed *in vivo* analyses have indicated that MreB structures turn over rapidly^10^. Purified MreB filaments are quite stable *in vitro*^11^, suggesting that MreB dynamics may be stimulated by accessory factors that have yet to be discovered.

For other well characterized cytoskeletal systems such as eukaryotic actin and tubulin and bacterial FtsZ, there are multiple known regulators of filament dynamics (reviewed in^12–14^). For MreB, in contrast, RodZ is the only confirmed regulator and it functions to stimulate MreB assembly^15^ and regulate filament properties^16^, leaving MreB disassembly mysterious. There are several toxin-antitoxin systems whose toxins have been proposed to target MreB^17–19^, but the degree to which these toxins are expressed and function under standard growth conditions remains unclear. The only other factor proposed to interact with MreB is MbiA, a small *C. crescentus* protein that interacts with MreB through an unknown mechanism^20^. The effects of MbiA on MreB also remain unclear, as MreB localization was characterized using a non-functional N-terminal fluorescent fusion to MreB^20^.

Here, we address the lack of knowledge of MreB assembly inhibitors by directly screening for such factors with an overexpression library. We chose an overexpression approach since MreB and many of its known interactors are essential. Our overexpression screen identified a new factor that we named <u>A</u>ssembly <u>I</u>nhibitor of <u>M</u>re<u>B</u> (AimB). As predicted for an important regulator of an essential gene, AimB appears to be essential. Overexpression of AimB resulted in wider cells that resemble the loss of MreB. To characterize the function of MreB, we developed a functional “sandwich” fusion of msfGFP to *C. crescentus* MreB and found that AimB inhibits its proper localization. Genetic and biochemical studies confirmed that AimB directly interacts with MreB. Finally, we used all-atom molecular dynamics simulations to develop a model for how AimB inhibits the assembly of MreB and confirmed predictions of this model biochemically.

## Results

### *A* C. crescentus *protein overexpression screen identifies a novel cytoskeletal regulator*

We previously constructed a *C. crescentus* Gateway entry vector library that includes 224 entry vectors containing ORFs encoding “conserved hypothetical” proteins^21^. To identify candidate MreB regulators among these previously uncharacterized proteins, we transferred these ORFs into a xylose-inducible overexpression destination vector using an *in vivo* Gateway cloning system, conjugated these constructs into *C. crescentus*, and imaged the strains at the single-cell level^21^. Among the various phenotypes observed, overexpression of *cc_2490* resulted in a significant increase in cell width that was similar to that seen upon disruption of MreB assembly by the small-molecule inhibitor A22^22^ (Figure 1A).

**Figure 1:**
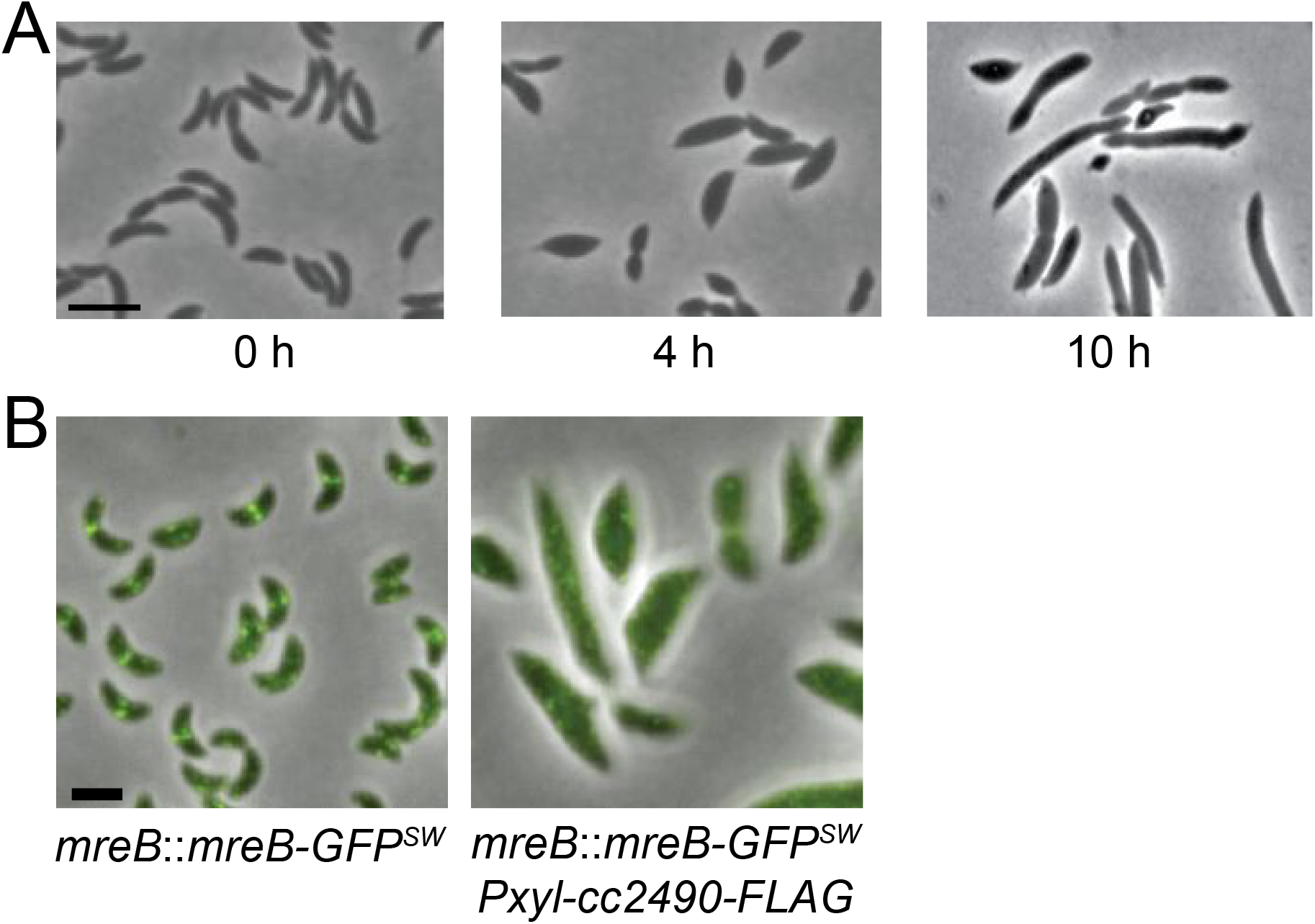
Cell width and MreB localization are disrupted by CC_2490 overexpression. A) CC_2490 expression was induced in wild-type *C. crescentus* for the indicated times. Phase-contrast images show disruption to cell width and cell shape in *C. crescentus* cells overexpressing CC_2490. Scale bar: 2 µm. B) MreB-GFP^sw^ cells with or without CC_2490 overexpression were imaged by phase and fluorescence microscopy at 9 h post-induction. MreB was delocalized in cells grown with CC_2490 overexpression. Scale bar: 2 µm.

We expected that a factor that disrupts MreB assembly would have a strong effect on MreB localization. Since previous analyses of MreB localization in *C. crescentus* used N-terminal fluorescent fusions that we now know to be non-functional^23^, we first developed a functional reporter for *C. crescentus* MreB localization. To this end, we inserted monomeric-superfolder GFP (msfGFP), which is less prone to aggregation than most commonly used fluorescent proteins, into the same surface-exposed loop that tolerates functional fusion insertions in *E. coli* MreB^24^. We replaced *mreB* at its native chromosomal locus under its native promoter to generate a strain in which the only copy of MreB is this new “sandwich” fusion (MreB-GFP^sw^). The MreB-GFP^sw^ fusion does not affect proliferation rate (Figure S1A), suggesting that it is functional with regards to regulating cell growth and division. As observed in the homologous msfGFP-fusion in *E. coli*, *C. crescentus* cells expressing MreB-GFP^sw^ were slightly wider and shorter than wild-type cells (Figure S1B,C).

Consistent with its effects on cell shape, overexpression of *cc_2490* strongly disrupted MreB localization. Whereas wild-type cells showed MreB-GFP^sw^ foci distributed in patches or at midcell in dividing cells, *cc_2490* overexpression caused MreB-GFP^sw^ to disperse and become diffuse or to accumulate at the poles (Figure 1B). Based on these morphological and MreB-localization phenotypes, we renamed CC_2490 AimB for <u>A</u>ssembly <u>I</u>nhibitor of <u>M</u>re<u>B</u>. AimB is a member of the Domain-of-Unknown-Function (DUF) superfamily DUF1476 and is widely conserved among Alphaproteobacteria but has no other known activity.

### AimB and A22 have additive effects

Since A22 treatment is lethal to *C. crescentus* cells, we examined whether AimB overexpression is toxic. After only a few hours of overexpression, we observed a significant drop in growth rate as measured by optical density and colony forming units, confirming that AimB overexpression is lethal (Figure 2A,B). Western blots for MreB showed no change in MreB protein levels when AimB is overexpressed (Figure S2A), demonstrating that AimB toxicity was not due to a reduction in MreB protein concentration. To compare the toxicity of AimB overexpression with that of A22, we measured growth with AimB overexpression and A22 treatment individually or in combination. Both treatments were toxic, and the combination of AimB overexpression and A22 treatment further enhanced lethality (Figure 2C,D).

**Figure 2:**
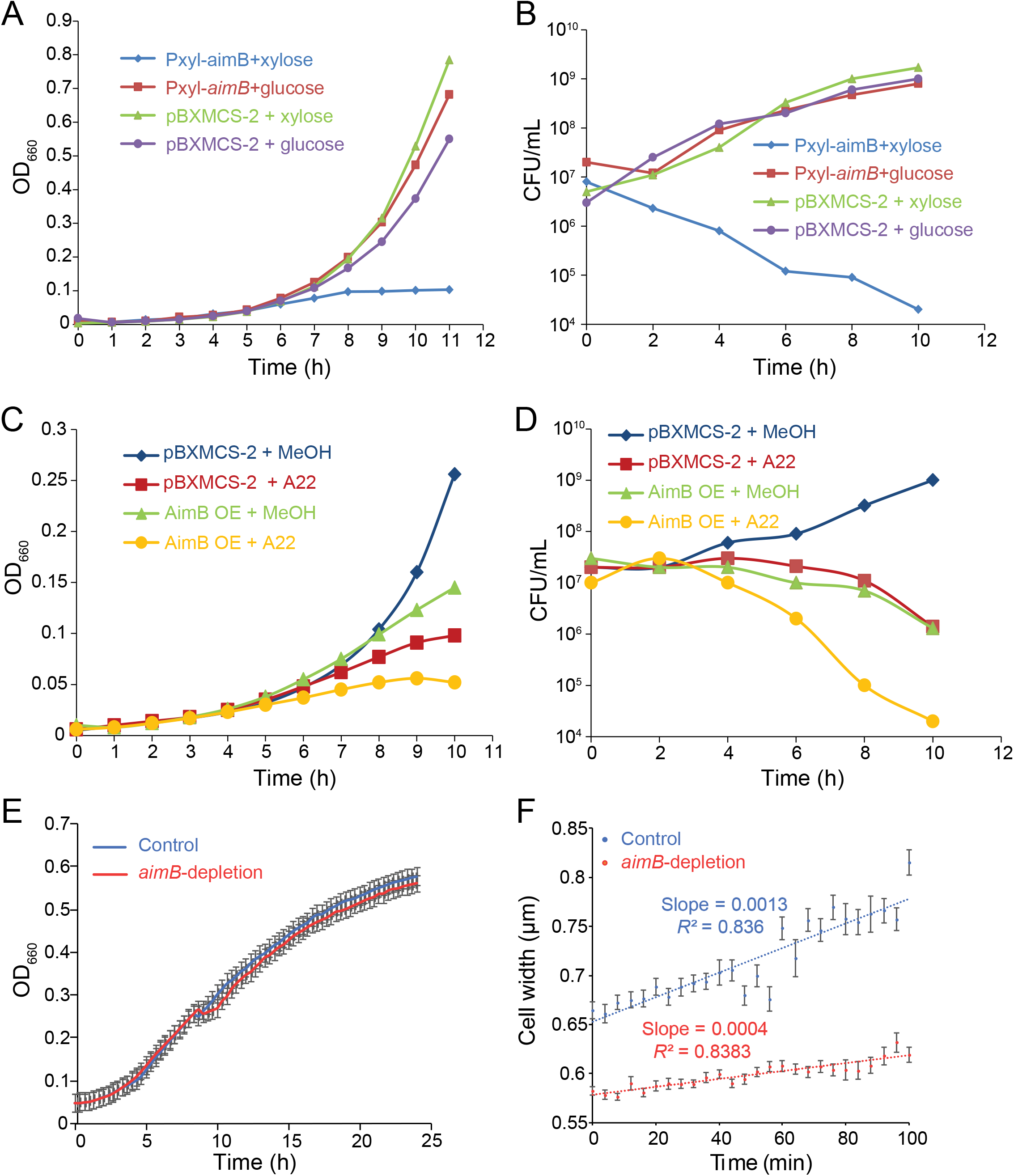
Regulation of AimB expression is critical for rapid *C. crescentus* growth. A) AimB overexpression inhibited growth as measured by optical density. Cells containing either the empty vector (pBXMCS-2) or an AimB overexpression vector were grown with 0.03% xylose (induced) or 0.3% glucose (uninduced). B) AimB overexpression was toxic to cells as measured by colony forming units (CFUs). Samples were removed every 2 h from the cultures in (A) and plated to measure CFUs. C,D) AimB overexpression and A22 treatment synergistically resulted in toxicity as measured by optical density (C) and CFUs (D). In (C), cells containing either pBXMCS-2 or an AimB overexpression vector were grown in the presence of 0.03% xylose and either 10 µg/mL A22 or methanol (MeOH). Samples were removed every 2 h from the cultures in (C) and plated to measure CFUs (D). E) Depletion of *aimB* mRNA using CRISPRi did not affect population growth as measured by optical density. dCas9 expression was induced with 0.5 mM vanillate in cells harboring either a control plasmid (psgRNA-base) or sgRNA-*aimB*. F) The rate of width increase for cells grown on PYE-agarose pads containing 2.5 µg/mL A22 and 0.5 mM vanillate was higher in cells depleted for AimB than in wild-type cells.

The similarities between A22 treatment and AimB overexpression suggested that AimB functions to destabilize MreB. Thus, we hypothesized that loss of AimB would stabilize MreB filaments. AimB appears to be essential for *C. crescentus* survival, as we were unable to generate a clean *aimB* deletion. Depletion of *aimB* using CRISPRi^25^ resulted in a 73.7 ± 2.0% (standard error of the mean; *n*=3) knockdown of *aimB* mRNA without having an effect on cell growth (Figure 2E). By contrast to the increased cell width upon AimB overexpression, depletion resulted in narrower cells when compared to controls (Figure 2F, initial time point). We hypothesize that the incomplete knockdown of *aimB* mRNA permitted cell proliferation despite having an effect on cell width. If loss of AimB stabilizes MreB filaments, we would expect these cells to have increased resistance to A22 treatment. Time-lapse imaging of control and *aimB*-depleted cells grown on A22-containing agarose pads demonstrated that the initial rate of cell-width increase was faster in the control cells (Figure 2F). Consistent with their opposing effects on MreB, *aimB* depletion doubled the minimum inhibitory concentration (MIC) of A22 compared to the control (control MIC = 2 µg/mL; *aimB*-depletion MIC = 4 µg/mL). Thus, AimB modulates MreB function in a manner consistent with that of a negative regulator of MreB assembly.

### AimB and MreB interact genetically

To identify the cellular targets of AimB, we performed a screen to identify suppressors of toxicity associated with AimB overexpression. Suppressors of overexpressed AimB-FLAG were isolated and subsequently screened by Western blot to filter out mutants with reduced AimB expression. This screening eliminated suppressors that decreased AimB production as well as nonsense and frameshift mutations in the *aimB* gene. For each isolated suppressor, we sequenced *aimB* from the overexpression vector and the chromosomal *mreB* gene. Three point mutations were identified in the overexpressed *aimB* that resulted in the residue changes V66M, L74Q, and A97P. Interestingly, 13 unique single point mutations were also found in *mreB*, demonstrating a genetic interaction between AimB and MreB.

To gain insight into the potential interaction between MreB and AimB we mapped the altered residues in MreB and AimB suppressors onto structures of *C. crescentus* MreB and an AimB homolog from *Jannaschia* sp. (Figure S3A-C). Two mutations, MreB^K236T^ and MreB^T277A^, were located at what is predicted to be the MreB longitudinal polymerization interface^26^. These changes may suppress the effects of AimB overexpression by stabilizing MreB filaments via increasing the interaction strength between MreB monomers or disrupting MreB-AimB interaction. The remaining 11 *mreB* mutations map near the ATP binding pocket and are reminiscent of mutations that suppress the effects of A22 treatment^27^.

Since similar *mreB* mutations can confer resistance to A22 treatment and AimB overexpression, we tested previously characterized strains with A22-resistant point mutations in *mreB*^27^ for their ability to suppress AimB overexpression. *C. crescentus* producing chromosomally-encoded MreB^T167A^, MreB^L23A^, MreB^D192G^, or MreB^V324A^ were resistant to AimB overexpression (Figure 3A). These mutations are predicted to inhibit ATP hydrolysis, thereby stabilizing MreB filaments. Conversely, bacteria expressing AimB-resistant *mreB* mutants also exhibited increased resistance to A22 (Figure 3B). Interestingly, the two MreB mutations at the longitudinal polymerization interface (K236T and T277A) had the highest sensitivity to 5 µg/mL A22 of all the mutants. This variability in A22 sensitivity demonstrates that mutations in the ATP binding pocket likely suppress the effects of AimB overexpression by a different mechanism than the mutations involved in MreB subunit-subunit interactions. Thus, while A22 treatment and AimB overexpression both appear to destabilize MreB polymers, they may act through distinct molecular mechanisms, which would explain their synergistic effects on cell growth.

**Figure 3:**
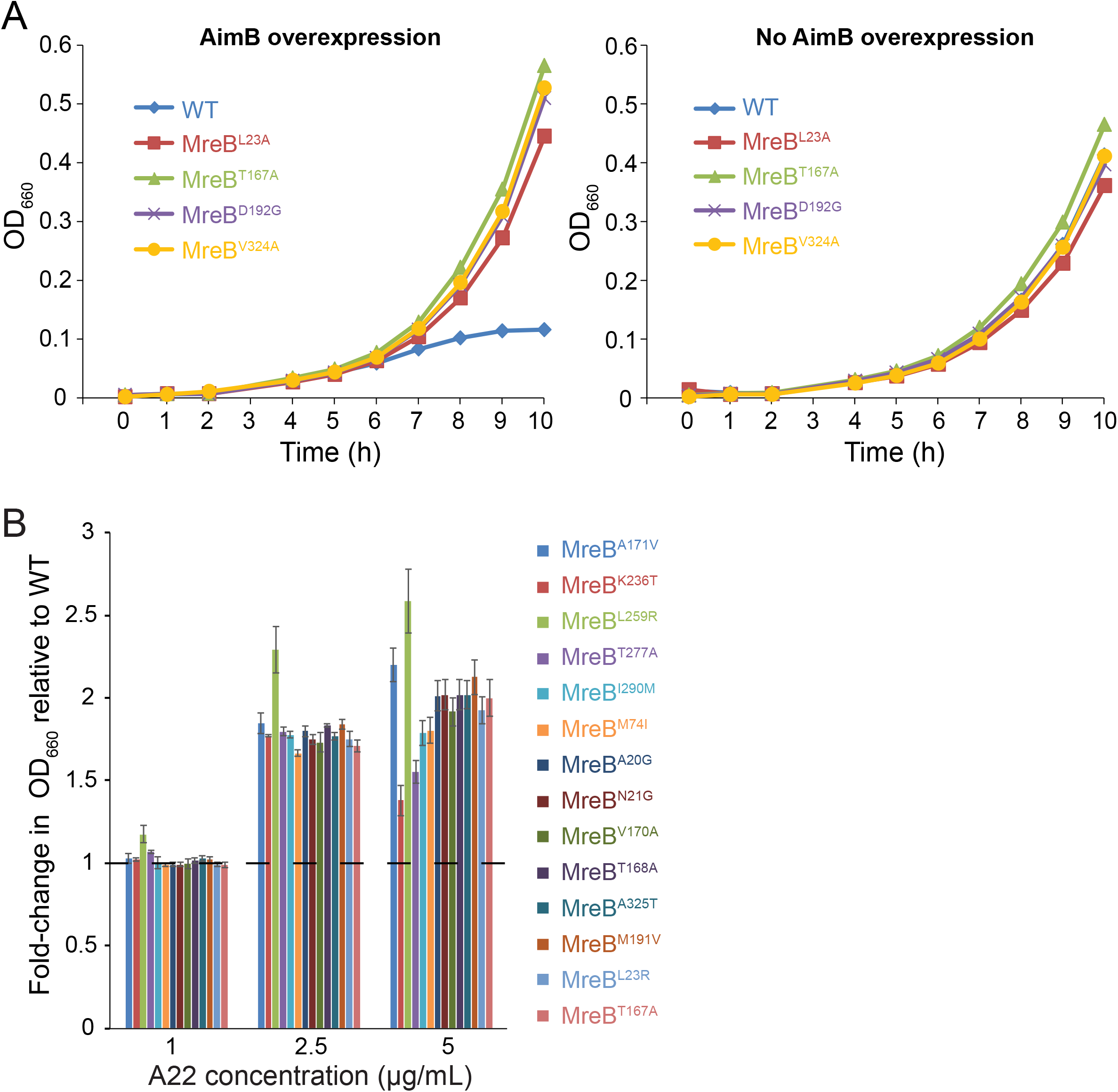
MreB mutations confer increased resistance to AimB overexpression and A22. A) A22 resistance mutations complemented the growth defect due to AimB overexpression. Growth curves for wild-type and A22-resistant strains containing an AimB overexpression plasmid grown with 0.03% xylose (AimB overexpression) or 0.3% glucose (no AimB overexpression). B) AimB overexpression-resistant strains exhibited higher resistance to A22 than wildtype. Overnight cultures were diluted and grown in media containing 1, 2.5, and 5 µg/mL A22 for 8 h, at which time OD_660_ readings were taken and standardized to the wild-type culture grown in that concentration of A22. Error bars are standard error of the mean (*n*=3).

### *The activity of AimB is specific to* C. crescentus *MreB*

AimB is highly conserved among Alphaproteobacteria but rarely found outside of this clade. Since AimB is essential in *C. crescentus* yet absent in *E. coli*, we tested whether AimB alters cell-shape and/or MreB localization in *E. coli*. Even when AimB was expressed at similar or slightly higher levels compared to those that have a strong impact in *C. crescentus* (Figure S2B), there was no effect on *E. coli* cell shape or on qualitative MreB localization (Figure 4A). This selectivity of AimB for *C. crescentus* MreB is particularly interesting given that the MreB orthologues in these organisms are 78% similar and 64% identical.

**Figure 4:**
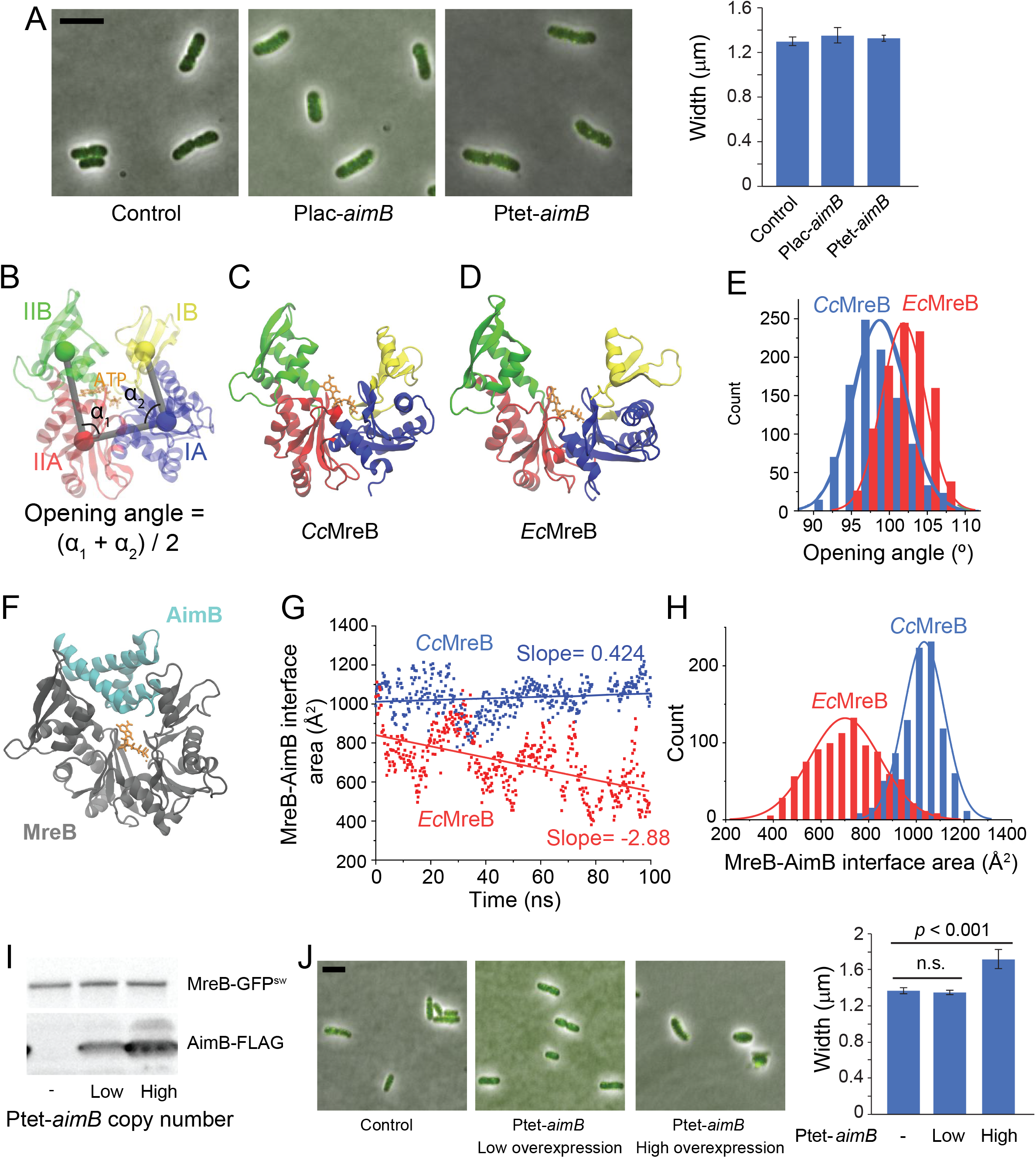
AimB has low affinity for *E. coli* MreB, potentially due to differences in the binding pocket. A) *E. coli* expressing MreB-GFP^sw^ were transformed with low-copy plasmids for AimB-FLAG expression and induced with 1 mM IPTG or 100 ng/mL aTc for 6 h. Cells were back-diluted 1:500 at 3 h to maintain log-phase growth. Cells were imaged by phase and fluorescence microscopy (overlay on left) and cell widths were analyzed (right). No effect on cell width or shape was observed. Scale bar = 5 µm. B) Definitions of opening angle for an MreB monomer. The centers-of-mass of the four subdomains are shown as colored spheres. C) Snapshot of an ATP-bound *Cc*MreB (PDB ID: 4CZM) at the end of a 100 ns simulation. D) Snapshot of an ATP-bound *Ec*MreB at the end of a 100 ns simulation demonstrating a larger opening angle than *Cc*MreB in (C). The initial *Ec*MreB structure was a homology model of *E. coli* MreB built from the *Cc*MreB crystal structure. E) *Ec*MreB exhibited larger opening angles than *Cc*MreB at the last 30 ns of MD simulations. F) Docking of a homology model of the *Jannaschia* sp. protein Jann_2546 (PDB ID: 2KZC), a homolog of AimB, to the equilibrated open structure from a *Cc*MreB MD simulation. G,H) The interfacial area between MreB and AimB showed that the docked heterodimer of *Cc*MreB and AimB in (F) remained stable throughout 100 ns of MD simulation (Movie S1), while the interfacial area of AimB docking to *Ec*MreB decreased over time (G). The distribution of interfacial areas over the course of the MD simulation demonstrates that AimB interacts more stably with *Cc*MreB (H). I,J) Substantial overexpression of AimB in *E. coli* disrupts cell width and MreB localization. *E. coli* MreB-GFP^sw^ strains harboring low-or high-copy vectors for AimB-FLAG expression were induced as in (A). In (I), cell lysates (normalized to OD_600_) were analyzed by immunoblotting. In (J), cellular dimensions were quantified by phase and fluorescence microscopy. Scale bar = 5 µm.

### Structural modeling suggests that AimB binds in the cleft of MreB

To develop a molecular hypothesis for how AimB specifically affects *C. crescentus* MreB, we used molecular dynamics (MD) simulations^28^ to investigate whether the differential effects of AimB in *E. coli* and *C. crescentus* are due to different conformations of the two proteins. We generated a homology model of *E. coli* MreB (*Ec*MreB) based on the *C. crescentus* MreB (*Cc*MreB) structure (PDB ID: 4CZM)^26^, and performed all-atom MD simulations (see Methods) on ATP-bound *Ec*MreB and *Cc*MreB monomers. We previously observed for *Thermatoga maritima* MreB monomers that the opening angle at the polymerization interface (Figure 4B) was polymerization dependent, with a larger value for monomers relative to the subunits of a dimer^28^. Here, we found that the opening angle of an *Ec*MreB monomer was significantly higher than that of *Cc*MreB (Figure 4C-E). Thus, we hypothesized that AimB’s selectivity could be explained by binding within the gap formed at the MreB-MreB longitudinal polymerization interface. Specifically, binding of AimB when *Cc*MreB monomers open would prevent binding by another MreB monomer and inhibit polymer assembly. Meanwhile, the larger opening in *Ec*MreB would destabilize the binding of AimB, rendering it less active.

In support of our hypothesis, we were able to dock an AimB homology model of the *Jannaschia* sp. protein Jann_2546 (PDB ID: 2KZC) to the equilibrated open structure of our *Cc*MreB MD simulations (Fig. 4F), and this docked heterodimer remained stable throughout 100 ns of MD simulation (Figure 4G and Movie S1). By contrast, after a docking of the *C. crescentus* AimB homology model to an *Ec*MreB with a similar opening angle to that of the equilibrated *Cc*MreB-ATP structure, the AimB gradually dissociated during the simulation (Figure 4G and Movie S2), coincident with further MreB opening. Quantification of the MreB-AimB interfacial area over 100 ns of simulation showed that AimB consistently had greater contact with *Cc*MreB as compared to *Ec*MreB (Figure 4H).

Our MD simulations suggested that AimB can form a stable interaction within the opening cleft at the longitudinal polymerization interface of *Cc*MreB, while AimB has decreased affinity for *Ec*MreB. Therefore, we hypothesized that the decreased affinity of AimB for *Ec*MreB could be overcome by increasing its expression. Consistent with this prediction, when we expressed *aimB* from a high-copy *E. coli* expression vector, we observed an increase in *E. coli* cell width (Figure 4I,J), similar to the effects of sublethal A22 treatment^29^. Importantly, the residues of *Cc*MreB that interact with AimB (within 5 Å) are highly conserved in *Ec*MreB (79% identical and 96% similar; Figure S3D); thus, the relative affinities for *Cc*MreB and *Ec*MreB appear to be due to their opening angles rather than differences in binding-site amino acids.

### AimB and MreB interact directly

The isolation of AimB-resistant strains with MreB mutations suggested that MreB and AimB interact directly, and our MD docking simulations further predicted specific regions of the two proteins that may interact. To test these predictions, we used a photo-crosslinking assay. Specifically, we created an expression plasmid with *C. crescentus* MreB driven by the *lac* promoter and AimB driven by an arabinose-inducible promoter. Based on the *Cc*MreB crystal structure, we selected 26 surface-accessible residues (Figure 5A) to probe for AimB interactions. Each of the 26 residues was individually mutated to the amber stop codon TAG to enable the incorporation of the unnatural amino acid p-benzoylphenylalanine (pBPA). Each amber mutant plasmid was transformed into an *E. coli* Δ*mreB* strain carrying the plasmid pEVOL-pBpF, which encodes the tRNA synthase/tRNA pair for pBPA incorporation^30^. We chose to use a Δ*mreB* strain so that the only potential MreB-AimB interaction would be that of the *C. crescentus* proteins. Following cross-linking, an interaction was only observed when pBPA was incorporated at residue 185 of MreB (Figure 5B). Probing this interaction with an anti-FLAG antibody to detect AimB-FLAG confirmed the interaction (Figure 5C). The size of the shifted band indicated a 1:1 interaction stoichiometry between MreB and AimB. Strikingly, this position is at the base of the cleft where AimB and MreB are predicted to interact based on MD simulations; analysis of our simulations showed that the intermolecular distance between MreB^R185^ and AimB^G64^ remained small in *Cc*MreB whereas the distance was larger and more variable in *Ec*MreB (Figure 5D). These crosslinking data provide compelling evidence that AimB directly interacts with MreB *in vivo* in a manner that validates the conclusions of our MD simulations.

**Figure 5:**
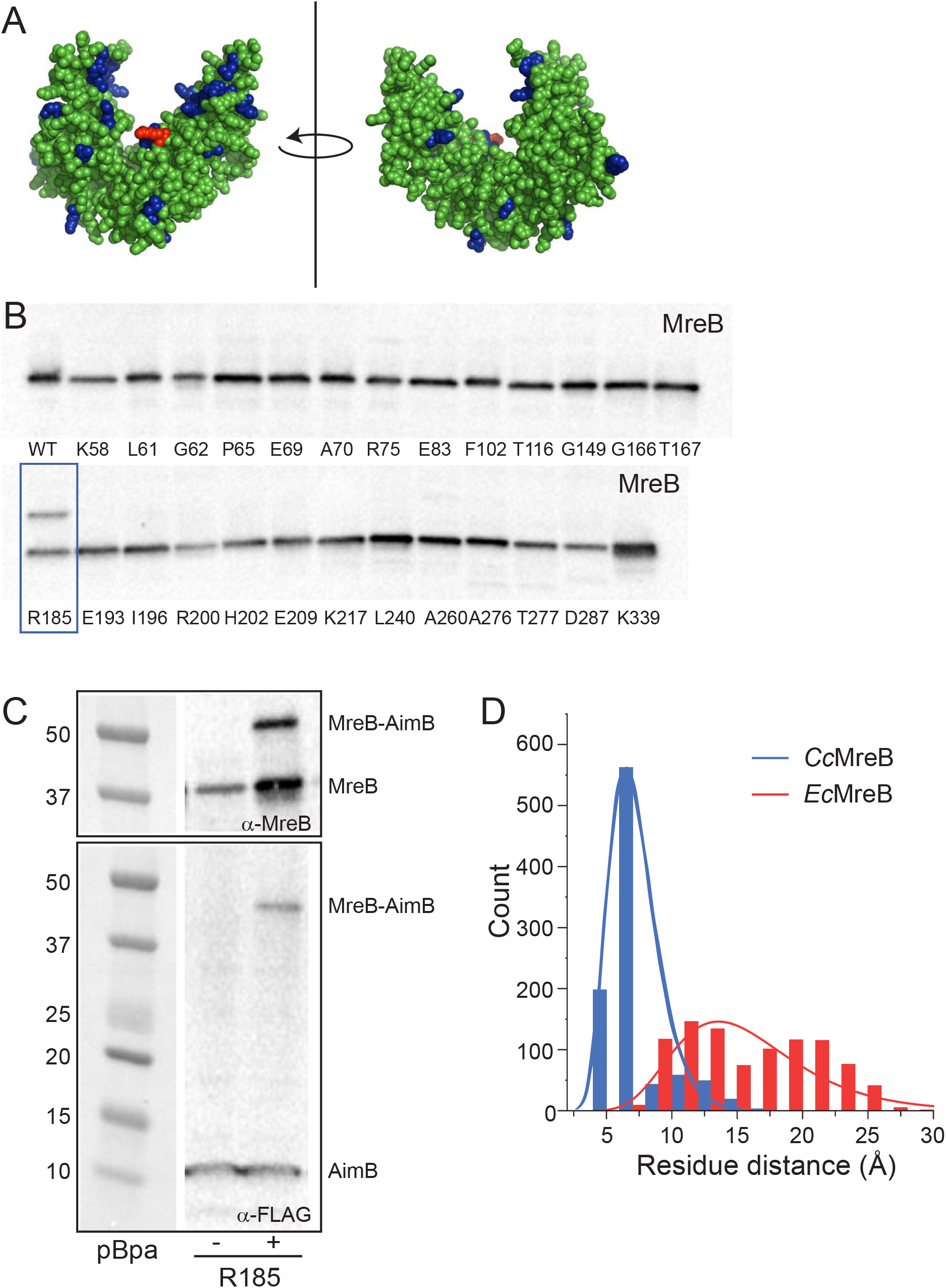
AimB and MreB interact directly. A) The location of the 26 residues of MreB that were mutated to the amber codon for *in vitro* crosslinking assays are highlighted in blue on the *Cc*MreB crystal structure. Arginine 185 is highlighted in red. B) *In vitro* crosslinking experiments were performed by incorporating the UV-crosslinkable non-natural amino acid p-benzoylphenylalanine at various positions in MreB (Materials and Methods). Crosslinked samples were analyzed by immunoblotting for MreB. A crosslinked band was observed for position R185 (blue rectangle). C) UV-crosslinking of R185 was performed as in (A) using a FLAG-tagged AimB construct. Immunoblotting for MreB or the FLAG-tagged AimB showed similar crosslinked bands. D) R185 is at the base of the cleft where AimB and MreB are predicted to interact. The intermolecular distance between MreB^R185^ and AimB^G64^, the nearest AimB residue, was quantified over the course of our *Cc*MreB-AimB and *Ec*MreB-AimB MD simulations. AimB interacts with *Cc*MreB more stably compared to *Ec*MreB, as shown by a smaller distance between the two residues.

## Discussion

As the number of sequenced bacterial genomes rapidly increases, a striking feature of virtually all genomes is the lack of comprehensive annotation, leading to an overwhelming number of “hypothetical genes” whose cellular functions are completely unknown. For even the best studied model organism, *E. coli* K-12, the fraction of hypothetical genes is >25% (UniProt “uncharacterized” or “putative” genes), roughly similar to other model organisms such as *Pseudomonas aeruginosa* PAO1 (39%), *Vibrio cholera* O1 El Tor (42%), *B. subtilis* 168 (43%), and *C. crescentus* (20%)^31^. Moreover, these fractions are likely an underestimate because automated genome annotation pipelines have difficulty distinguishing bona fide small proteins from random unexpressed open reading frames. Advanced transcriptomics and proteomics techniques, such as ribosome profiling^32^, have improved our ability to robustly confirm the expression of small proteins (<50 residues), some of which are critical regulators of protein kinases, membrane bound enzymes, transport, cell division, or spore formation (reviewed in^33^). Using the *C. crescentus* genome as an example, of the 762 genes annotated as hypothetical proteins, there are 34 ORFs shorter than 50 codons and 172 ORFs with 50-100 residues. Here we establish overexpression phenotypic screening as a rapid and robust platform to functionally characterize hypothetical proteins involved in the regulation of the bacterial cytoskeletal element MreB.

The turnover of eukaryotic actin filaments is accomplished by a variety of regulatory proteins that either sequester actin monomers or sever intact filaments^34,35^. While structural studies of actin-regulator interactions have yielded mechanistic insights into the modulation of actin polymerization, our understanding of MreB polymerization dynamics in general and polymer turnover in particular is quite limited. In *C. crescentus*, the protein MbiA binds directly to MreB, and its overexpression leads to a loss of proper cell shape and an increase in cell death^20^. The *E. coli* toxins YeeV and CptA inhibit MreB polymerization *in vitro*^18,19^, however their roles in normal physiology are unclear. Importantly, the mechanism of action for all three proposed MreB inhibitors is unknown. Here, we identified AimB as a novel inhibitor of *C. crescentus* MreB and provide the first mechanistic model for MreB assembly inhibition. Specifically, *in vivo* cross-linking experiments (Figure 5A-D) coupled with MD simulations (Figure 4B-H) suggest a novel mechanism for the interaction between AimB and MreB in the cleft of open MreB subunits that blocks MreB dimerization.

Overexpression of AimB results in an increase in cell width (Figure 1A) and mislocalization of MreB (Figure 1B) in a manner similar to the MreB inhibitor A22. A screen for AimB-overexpression suppressor mutants found mutations in MreB (Figure 3B), demonstrating a genetic interaction between MreB and AimB. To probe for a direct interaction between these proteins, we used a photo-crosslinking approach to discover that MreB residue 185 interacts with AimB (Figure 5B,C). These data are consistent with our MD simulations that propose a model in which AimB binds to the longitudinal polymerization interface of MreB. In this model, AimB would function as a pointed-end competitive inhibitor of MreB-MreB dimerization. This model represents a novel mechanism for destabilizing actin-like filaments; thymosin-β4 sequesters G-actin monomers by stretching across the actin molecule and interacting with both the pointed and barbed ends^36^, while twinfilin inhibits actin polymerization by binding G-actin barbed ends with high affinity^37^.

In addition to explaining how AimB inhibits MreB assembly, our model can also explain the specificity of AimB for *C. crescentus* MreB as well as the synergy between AimB overexpression and A22 treatment. Our simulations and crosslinking are consistent with AimB binding to the cleft that forms in MreB subunits at the longitudinal (intra-polymeric) polymer interface when the opening angle is large. Binding of AimB at this site would sterically prevent additional MreB monomers from adding to the polymer, thereby inhibiting MreB filament assembly. Furthermore, this binding site is conformationally distinct in *C. crescentus* and *E. coli* (Figure 4B-E), thereby explaining the species specificity. Finally, the structure of *C. crescentus* MreB filaments solved in the presence of A22 indicates that A22 disrupts the lateral (inter-polymeric) MreB filament interface^26^, which would explain how AimB and A22 use distinct mechanisms to inhibit MreB filament formation and therefore synergistically inhibit growth rate.

MreB coordinates peptidoglycan insertion to regulate cellular elongation in a variety of species, including Gram-negative *E. coli*^7^ and Gram-positive *B. subtilis*^5,6^. Although *mreB* is found across a wide range of bacterial lineages, the *aimB* gene is restricted to Alphaproteobacteria. Based on our overexpression studies and MD simulations, we suggest that AimB binds the longitudinal polymerization interface of *C. crescentus* MreB with higher affinity than *E. coli* MreB. This species specificity is demonstrated by the ability of AimB to disrupt the localization of *E. coli* MreB only when highly overexpressed (Figure 4J). Species-specific regulation of a bacterial cytoskeletal protein is not unexpected given that the highly conserved tubulin-ortholog FtsZ is regulated by a variety of divergent mechanisms. For example, placement of FtsZ and the divisome at midcell can be mediated by multiple, distinct mechanisms. In *E. coli* and most Gram-negative bacteria, oscillations of the MinC/D complex are facilitated by MinE^38,39^, whereas in *B. subtilis* and most Gram-positive bacteria MinC/D restricts FtsZ to the midline via interactions with DivIVA^40^. Similarly, nucleoid occlusion in *E. coli* and *B. subtilis* is directed by two different proteins, Noc and SlmA, respectively^41,42^. Interestingly, *C. crescentus* does not use either the MinC/D or nucleoid occlusion mechanisms for FtsZ localization; instead, a gradient of MipZ antagonizes FtsZ polymerization closer to the poles, leading to midcell Z-ring formation^43^. Thus, while the core MreB and FtsZ cytoskeletal proteins are widely conserved in bacteria, emerging evidence suggests that the regulation of these core cytoskeletons is largely performed by species-specific factors.

## Methods

### Bacterial strains, plasmids, and growth conditions

The strains, plasmids, and primers used in this study are described in Tables S1, S2, and S3, respectively. Details regarding strain construction are available in the Supplementary Text. *C. crescentus* wild-type strain CB15N and its derivatives were grown at 30 °C in peptone-yeast-extract (PYE) medium (Poindexter, 1964). *E. coli* strains were grown at 37 °C in LB medium. When necessary, antibiotics were added at the following concentrations: kanamycin (Kan) 30 µg/mL in broth and 50 µg/mL in agar (abbreviated 30:50) for *E. coli* and 5:25 for *Caulobacter*; tetracycline (Tet) 1:2 for *Caulobacter*; chloramphenicol (Cm) 20:30 for *E. coli*; carbenicillin (Carb) 50:100 for *E. coli*. Gene expression was induced in *Caulobacter* (0.03-0.3% w/v xylose; 0.5 mM vanillate) or *E. coli* (100 ng/mL anhydro-tetracycline (aTc)); 1 mM isopropyl β-D-1-thiogalactopyranoside (IPTG)) as noted. Pharmacological inhibition of MreB was performed by adding 1-10 µg/mL A22 (methanol was used as the vehicle control).

### CRISPRi-mediated gene depletion

*C. crescentus* CRISPRi was performed using the plasmids (Table S2) and methods developed by the Jacobs-Wagner lab^25^. Briefly, primers EK1003/1004 (Table S3), encoding the sgRNA mapping to the 5’-end of *aimB*, were phosphorylated and annealed. The annealed oligos were ligated into the BbsI site of plasmid psgRNA-Base. The resulting plasmid (pEK334) was transformed into a strain carrying a vanillate-inducible catalytically dead *cas9* gene (CJW6270) to generate strain EK335 (Δ*vanA*::pV-dcas9hum-RBSmut1 with plasmid psgRNA-*aimB*). Gene-depletion was initiated with 0.5 mM vanillate and monitored by qRT-PCR. Cells carrying psgRNA-base were used as controls.

### High-throughput cloning and microscopy

Xylose-inducible plasmids for overexpression of conserved hypothetical proteins were generated using an *in vivo* Gateway strategy, as described previously^21,44^. The resulting multicopy plasmids were conjugated into *C. crescentus*. Strains were induced with 0.3% xylose and imaged in high-throughput format using custom 48-pedestal agarose slides^21,44^. Cell morphology was compared to wild-type controls to identify overexpression plasmids resulting in aberrant cell shape.

### Fluorescence microscopy and image analysis

Cells were spotted onto pads made of 1% agarose with the corresponding growth medium. Fluorescence microscopy was performed on a Nikon Ti-E inverted microscope equipped with a Lumen 220PRO illumination system (Prior), Zyla sCMOS 5.5-megapixel camera (Andor), CFI Plan Apochromat 100X oil immersion objective (NA 1.45, WD 0.13 mm), and NIS Elements software for image acquisition. Images were segmented using *Morphometrics*^45^. Cell width and length were calculated using custom Matlab scripts. For time-lapse imaging, coverslips were sealed with VALAP (1:1:1 vaseline:lanolin:paraffin) to prevent drying of the agarose pad.

### Cell growth measurements

For experiments up to 12 h, cells were grown in standard culture tubes and aliquots were removed at the specified intervals for measurements of OD660 or colony forming units (CFUs). For experiments longer than 12 h, cells were aliquoted into a 96-well plate and the OD660 was measured on a ClarioSTAR plate reader (BMG Labtech) with shaking and temperature control.

### Immunoblotting

Cell samples were normalized by optical density (1 mL of OD=0.5) and lysed in 1X SDS sample buffer. Samples were separated on a 4-20% gradient polyacrylamide gel, transferred to a PVDF membrane, and blotted with antibodies against MreB (1:1000)^22^, GFP (1:1000, Abcam ab6556), or FLAG (1:500, Santa Cruz sc-166355). Horseradish peroxidase-conjugated secondary antibodies (1:5000) and enhanced chemiluminescence reagents (GE Healthcare) were used to detect the bands on a Bio-Rad ChemiDoc MP system.

### Quantitative RT-PCR (qRT-PCR)

RNA was extracted from bacterial cultures using the Qiagen RNeasy kit. Following DNase digestion, RNA (5 ng/µL) was reverse-transcribed using the High Capacity cDNA Reverse Transcription Kit (Applied Biosystems). 1 µL of cDNA was used as template in a 10 µL qRT-PCR reaction performed with Power SYBR reagent (Applied Biosystems). qRT-PCR was performed on an ABI QuantStudio 6 using the ∆∆Ct method. *rpoD* expression was used as the loading control.

### Molecular dynamics simulations

All simulations (Table S4) were performed using the molecular dynamics package NAMD v. 2.10^46^ with the CHARMM27 force field, including CMAP corrections^47^. Water molecules were described with the TIP3P model^48^. Long-range electrostatic forces were evaluated by means of the particle-mesh Ewald summation approach with a grid spacing of <1 Å. An integration time step of 2 fs was used^49^. Bonded terms and short-range, nonbonded terms were evaluated every time step, and long-range electrostatics were evaluated every other time step. Constant temperature (*T* = 310 K) was maintained using Langevin dynamics^50^, with a damping coefficient of 1.0 ps^−1^. A constant pressure of 1 atm was enforced using the Langevin piston algorithm^51^ with a decay period of 200 fs and a time constant of 50 fs. Setup, analysis, and rendering of the simulation systems were performed with the software VMD v. 1.9.2^52^.

### Simulated systems

MD simulations performed in this study are described in Table S4. Simulations were initialized from the *C. crescentus* MreB crystal structure (PDB ID: 4CZM)^26^. The bound nucleotide was replaced by ATP, and Mg^2+^-chelating ions were added for stability. An AimB homology model was built based on *Jannaschia sp.* protein Jann_2546 (PDB ID: 2KZC) using Phyre2^53^. Water and neutralizing ions were added around each monomer or dimer, resulting in final simulation sizes of up to 89,000 atoms. All simulations were run for 100 ns. For mean values and distributions of measurements, only the last 30 ns were used. To ensure simulations had reached equilibrium, measurement distributions were fit to a Gaussian.

### Analysis of opening angles

The centers-of-mass of the four subdomains of each protein were obtained using VMD. For each time step, we calculated one opening angle from the dot product between the vector defined by the centers-of-mass of subdomains IIA and IIB and the vector defined by the centers-of-mass of subdomains IIA and IA. Similarly, we calculated a second opening angle from the dot products between the vectors defined by the centers-of-mass of subdomains IA and IB and of subdomains IA and IIA. The opening angles we report are the average of these two opening angles. Subdomain definitions are as in^28^.

### *In vitro* crosslinking

A low-copy plasmid for induction of MreB and AimB was constructed using the pZS2-123 vector backbone^54^. The aTc-regulated CFP open reading frame was removed by inverse-PCR with primers EK644 and EK645 (Table S3). The arabinose-inducible RFP was replaced with AimB by Gibson assembly (primers EK646-649; Table S3). Wild-type MreB and a series of amber codon mutants (Table S1) were synthesized by Genscript (Piscataway, NJ) and used to replace the IPTG-inducible YFP to create a plasmid encoding P*lac*-MreB and P*ara*-AiMB (pMreBXL1-26). A C-terminal FLAG tag was introduced into AimB in a subset of amber codon mutants by inverse PCR using primers EK679-680. *In vitro* crosslinking of MreB and AimB was performed essentially as previously described^30^. pMreBXL and pEVOL-pBpF^30^ were co-transformed into strain NO36 (MC4100 ∆*mreB*) and grown overnight in LB containing kanamycin and chloramphenicol. Cells were diluted 1:100 into fresh LB with antibiotics along with inducers (1 mM IPTG and 0.1% w/v L(+)-arabinose) and 1 mM p-benzoylphenylalanine (Bachem). After 4 h, 1 mL of each culture was pelleted, resuspended in 50 µL cold PBS, and transferred to a white 96-well plate. The samples were irradiated under a UV bulb (Norman Lamps CFL15/UV/MED) on ice for 15 min and 50 µL 2X SDS sample buffer was added to stop the reaction. Samples were boiled for 5 min and analyzed by immunoblotting.

## Acknowledgements

We thank Anna Konovalova (University of Texas, Health Science Center) for helpful discussions and assistance with the photo-crosslinking assay and Christine Jacobs-Wager (Yale University) for providing CRISPRi reagents. Funding was provided by NSF CAREER Award MCB-1149328, the Stanford Center for Systems Biology under Grant P50-GM107615, and the Allen Discovery Center at Stanford on Systems Modeling of Infection (to K.C.H.); NIH Ruth L. Kirschstein National Research Service Award 1F32GM100677 (to J.H.); an Agilent Fellowship and a Stanford Interdisciplinary Graduate Fellowship (to H.S.); NIH Grant R01GM107384 (to Z.G.); and NSF CAREER Award MCB-1553004 (to E.A.K.). K.C.H. is a Chan Zuckerberg Investigator. All simulations were performed with computer time provided by the Extreme Science and Engineering Discovery Environment (XSEDE), which is supported by National Science Foundation grant number OCI-1053575, with allocation number TG-MCB110056 (to K.C.H.).

## Author contributions

J.N.W., H.S., J.H., K.C.H., Z.G., and E.A.K. designed the research, J.N.W., H.S., J.H., and E.A.K. performed research, and J.N.W., H.S., J.H., K.C.H., Z.G., and E.A.K. analyzed data. E.A.K. wrote the manuscript and J.N.W., H.S., K.C.H., Z.G., and E.A.K. edited the manuscript.

