## Supplementary material for "AimB is a small protein regulator of cell size and MreB assembly": Figure S

#### Supplementary methods

#### Strain construction

<u>Xylose-inducible expression of conserved hypothetical proteins.</u> The cloning of the entry-vector set of *C. crescentus* open reading frames (ORFs) was described previously[1]. Plasmid pZG241, which contains a xylose-inducible promoter upstream of a Gateway cloning cassette, was used as the destination vector[2]. The 224 ORFs corresponding to conserved hypothetical proteins were inserted into pZG241 using a previously described *in vivo* LR reaction[1,3].

<u>Chromosomal expression of *mreB-GFP*$^{SW}$.</u> The gene encoding msfGFP was cloned at the site of a loop-domain of MreB using Gibson assembly. The pNPTS-138 plasmid backbone was PCR amplified using primers pNTPS-F/R, the *mreB* upstream and downstream chromosomal DNA fragments were amplified using primers MreB-up/down-F/R, and msfGFP was amplified using msfGFP-F/R. The four fragments were joined using Gibson assembly. The resulting plasmid (pZG1534) was transformed into *C. crescentus*. Individual colonies were grown overnight in PYE without antibiotic selection and streaked onto PYE-3% sucrose plates to select for bacteria that lost the *sacB* cassette. Individual colonies were screened for MreB fluorescence (Strain ZG1511).

<u>AimB overexpression strains.</u> To generate *C. crescentus* AimB overexpression strains, the *aimB* gene was amplified using primers aimB-F/R and inserted into the NdeI/EcoRI site of pBXMCS-2 (plasmid pZG825). A FLAG-tag was added to the C-terminus of AimB by inverse-PCR with primers aimB-FLAG-F/R (plasmid pZG826). These plasmids were electroporated into NA1000

or *mreB::mreB-GFP*$^{SW}$ to generate strains ZG870, ZG871, and EK25. To generate low-copy *E. coli* AimB overexpression strains, *aimB-FLAG* was PCR amplified from pZG826 using primers EK809/810 (Ptet) or EK813/814 (Plac). Plasmid backbones were amplified with primers EK807/808 (pBbS2k, Ptet) or EK811/812 (pTrc99a, Plac). The resulting PCR fragments were ligated by Gibson assembly yielding plasmids pEK188 (Plac) and pEK189 (Ptet); the plasmids were transformed into ZG1516 (*mreB::mreB-GFP*$^{SW}$) to generate strains EK191 and EK192. For high-copy expression, Ptet-*aimB-FLAG* was amplified from pEK189 using primers EK840/841 and plasmid pUC19 was amplified with primers EK838/839. The PCR products were ligated by Gibson assembly and the resulting plasmid (pEK199) was transformed into ZG1516 to yield strain EK200.

**Table S1. Strains used in this study.**

| *C. crescentus* | | | |
|---|---|---|---|
| **Strain** | **Genotype** | **Construction** | **Source** |
| NA1000 | Synchronizable variant of wild-type *C. crescentus* strain CB15 | | [4] |
| ZG949 | Pxyl-*aimB* (Gateway) | *In vivo* Gateway cloning of conserved hypothetical genes | This study |
| ZG1511 | *mreB::mreB-GFP*$^{SW}$ | Transformation of NA1000 with pZG1534 followed by sucrose selection | This study |
| ZG870 | pBXMCS-Pxyl-*aimB* | Transformation of NA1000 with pZG825 | This study |
| ZG871 | pBXMCS-Pxyl-*aimB*-FLAG | Transformation of NA1000 with pZG826 | This study |
| EK25 | *mreB::mreB-GFP*$^{SW}$; pBXMCS-Pxyl-*aimB*-FLAG | Transformation of ZG1511 with pZG826 | This study |
| ZG883 | pBXMCS-2 | Transformation of NA1000 with pBXMCS-2 | This study |
| CJW5939 | Δ*vanA::*pV-dcas9hum-RBSmut1 with plasmid psgRNA-base | | [5] |
| EK335 | Δ*vanA::*pV-dcas9hum-RBSmut1 with plasmid psgRNA-*aimB* | Transformation of CJW6270 with pEK334 | This study |
| CJW6270 | Δ*vanA::*pV-dcas9hum-RBSmut1 | | [5] |
| JAT684 | *PmreB::T167A-mreB* | | [6] |
| JAT669 | *PmreB::L23P-mreB* | | [6] |
| JAT692 | *PmreB::D192G-mreB* | | [6] |
| JAT699 | *PmreB::V324A-mreB* | | [6] |
| ZG896 | *PmreB::L23P-mreB*; pBXMCS-AimB | Transformation of JAT669 with pZG825 | This study |
| ZG897 | *PmreB::T167A-mreB*; pBXMCS-AimB | Transformation of JAT684 with pZG825 | This study |
| ZG898 | *PmreB::D192G-mreB*; pBXMCS-AimB | Transformation of JAT692 with pZG825 | This study |
| ZG899 | *PmreB::V324A-mreB*; pBXMCS-AimB | Transformation of JAT699 with pZG825 | This study |
| ZG917 | *PmreB::A171V-mreB* | AimB-overexpression suppressor screen | This study |
| ZG918 | *PmreB::K236T-mreB* | AimB-overexpression suppressor screen | This study |

| | | | |
|---|---|---|---|
| ZG920 | *PmreB::T277A-mreB* | AimB-overexpression suppressor screen | This study |
| ZG921 | *PmreB::I290M-mreB* | AimB-overexpression suppressor screen | This study |
| ZG922 | *PmreB::M74I-mreB* | AimB-overexpression suppressor screen | This study |
| ZG923 | *PmreB::A20G-mreB* | AimB-overexpression suppressor screen | This study |
| ZG924 | *PmreB::N21G-mreB* | AimB-overexpression suppressor screen | This study |
| ZG925 | *PmreB::V170A-mreB* | AimB-overexpression suppressor screen | This study |
| ZG926 | *PmreB::T168A-mreB* | AimB-overexpression suppressor screen | This study |
| ZG928 | *PmreB::A325T-mreB* | AimB-overexpression suppressor screen | This study |

| *E. coli* | | | |
|---|---|---|---|
| **Strain** | **Genotype** | **Construction** | **Source** |
| S17-1 | λ–pir cloning strain, $Spec^R$ | | [7] |
| XL1-Blue | Cloning strain, $Tet^R$ | | Agilent Technologies |
| ZG1516 | *mreB::mreB-GFP*$^{SW}$ | | [8] |
| EK191 | *mreB::mreB-GFP*$^{SW}$; Plac-*aimB*-FLAG | Transformation of ZG1516 with pEK188 | This study |
| EK192 | *mreB::mreB-GFP*$^{SW}$; Ptet-*aimB*-FLAG (low copy) | Transformation of ZG1516 with pEK189 | This study |
| EK200 | *mreB::mreB-GFP*$^{SW}$; Ptet-*aimB*-FLAG (high copy) | Transformation of ZG1516 with pEK199 | This study |
| NO36 | Δ*mreB* (MC4100) | | Lab collection |
| EK85 | Δ*mreB*; pEVOL-pBpF | Transformation of NO36 with pEVOL-pBpF | This study |
| MreBXL-con | Δ*mreB*; pEVOL-pBpF; Plac-*mreB*/Para-*aimB* | Transformation of EK85 with pMreBXL-con | This study |
| MreBXL-1 | Δ*mreB*; pEVOL-pBpF; Plac-*mreB*$^{K58}$/Para-*aimB* | Transformation of EK85 with pMreBXL-1 | This study |
| MreBXL-2 | Δ*mreB*; pEVOL-pBpF; Plac-*mreB*$^{L61}$/Para-*aimB* | Transformation of EK85 with pMreBXL-2 | This study |
| MreBXL-3 | Δ*mreB*; pEVOL-pBpF; Plac-*mreB*$^{G62}$/Para-*aimB* | Transformation of EK85 with pMreBXL-3 | This study |
| MreBXL-4 | Δ*mreB*; pEVOL-pBpF; Plac-*mreB*$^{P65}$/Para-*aimB* | Transformation of EK85 with pMreBXL-4 | This study |
| MreBXL-5 | Δ*mreB*; pEVOL-pBpF; Plac-*mreB*$^{E69}$/Para-*aimB* | Transformation of EK85 with pMreBXL-5 | This study |

| | | | |
|---|---|---|---|
| MreBXL-6 | Δ*mreB*; pEVOL-pBpF; Plac-*mreB*$^{A70}$/Para-*aimB* | Transformation of EK85 with pMreBXL-6 | This study |
| MreBXL-7 | Δ*mreB*; pEVOL-pBpF; Plac-*mreB*$^{R75}$/Para-*aimB* | Transformation of EK85 with pMreBXL-7 | This study |
| MreBXL-8 | Δ*mreB*; pEVOL-pBpF; Plac-*mreB*$^{E83}$/Para-*aimB* | Transformation of EK85 with pMreBXL-8 | This study |
| MreBXL-9 | Δ*mreB*; pEVOL-pBpF; Plac-*mreB*$^{F102}$/Para-*aimB* | Transformation of EK85 with pMreBXL-9 | This study |
| MreBXL-10 | Δ*mreB*; pEVOL-pBpF; Plac-*mreB*$^{T116}$/Para-*aimB* | Transformation of EK85 with pMreBXL-10 | This study |
| MreBXL-11 | Δ*mreB*; pEVOL-pBpF; Plac-*mreB*$^{G149}$/Para-*aimB* | Transformation of EK85 with pMreBXL-11 | This study |
| MreBXL-12 | Δ*mreB*; pEVOL-pBpF; Plac-*mreB*$^{G166}$/Para-*aimB* | Transformation of EK85 with pMreBXL-12 | This study |
| MreBXL-13 | Δ*mreB*; pEVOL-pBpF; Plac-*mreB*$^{T167}$/Para-*aimB* | Transformation of EK85 with pMreBXL-13 | This study |
| MreBXL-14 | Δ*mreB*; pEVOL-pBpF; Plac-*mreB*$^{R185}$/Para-*aimB* | Transformation of EK85 with pMreBXL-14 | This study |
| MreBXL-15 | Δ*mreB*; pEVOL-pBpF; Plac-*mreB*$^{E193}$/Para-*aimB* | Transformation of EK85 with pMreBXL-15 | This study |
| MreBXL-16 | Δ*mreB*; pEVOL-pBpF; Plac-*mreB*$^{I196}$/Para-*aimB* | Transformation of EK85 with pMreBXL-16 | This study |
| MreBXL-17 | Δ*mreB*; pEVOL-pBpF; Plac-*mreB*$^{R200}$/Para-*aimB* | Transformation of EK85 with pMreBXL-17 | This study |
| MreBXL-18 | Δ*mreB*; pEVOL-pBpF; Plac-*mreB*$^{H202}$/Para-*aimB* | Transformation of EK85 with pMreBXL-18 | This study |
| MreBXL-19 | Δ*mreB*; pEVOL-pBpF; Plac-*mreB*$^{E209}$/Para-*aimB* | Transformation of EK85 with pMreBXL-19 | This study |
| MreBXL-20 | Δ*mreB*; pEVOL-pBpF; Plac-*mreB*$^{K217}$/Para-*aimB* | Transformation of EK85 with pMreBXL-20 | This study |
| MreBXL-21 | Δ*mreB*; pEVOL-pBpF; Plac-*mreB*$^{L240}$/Para-*aimB* | Transformation of EK85 with pMreBXL-21 | This study |
| MreBXL-22 | Δ*mreB*; pEVOL-pBpF; Plac-*mreB*$^{A260}$/Para-*aimB* | Transformation of EK85 with pMreBXL-22 | This study |
| MreBXL-23 | Δ*mreB*; pEVOL-pBpF; Plac-*mreB*$^{A276}$/Para-*aimB* | Transformation of EK85 with pMreBXL-23 | This study |
| MreBXL-24 | Δ*mreB*; pEVOL-pBpF; Plac-*mreB*$^{T277}$/Para-*aimB* | Transformation of EK85 with pMreBXL-24 | This study |
| MreBXL-25 | Δ*mreB*; pEVOL-pBpF; Plac-*mreB*$^{D287}$/Para-*aimB* | Transformation of EK85 with pMreBXL-25 | This study |
| MreBXL-26 | Δ*mreB*; pEVOL-pBpF; Plac-*mreB*$^{K339}$/Para-*aimB* | Transformation of EK85 with pMreBXL-26 | This study |
| EK210 | Δ*mreB*; pEVOL-pBpF; Plac-*mreB*$^{R185}$/Para-*aimB*-FLAG | Transformation of EK85 with pMreBXL-14-FLAG | This study |

**Table S2. Plasmids used in this study.**

| Name | Description | Source |
|---|---|---|
| pZG241 | Gateway destination vector based on pJS14 modified to include a xylose-inducible promoter, $Tet^R$ | [2] |
| pNPTS-138 | *sacB* expressing plasmid used for allelic exchange in *C. crescentus*, $Kan^R$ | M.R.K. Alley, unpublished |
| pZG1534 | pNPTS-138 based plasmid for replacing the chromosomal *C. crescentus mreB* with *mreB-GFP*$^{SW}$, $Kan^R$ | This study |
| pBXMCS-2 | *C. crescentus* high-copy plasmid with a xylose-inducible promoter, $Kan^R$ | [9] |
| pZG825 | pBXMCS-2 based plasmid for *aimB* expression, $Kan^R$ | This study |
| pZG826 | pBXMCS-2 based plasmid for *aimB-FLAG* expression, $Kan^R$ | This study |
| psgRNA-Base | sgRNA cloning vector for CRISPRi, $Kan^R$ | [5] |
| pEK334 | *aimB* sgRNA expression plasmid, $Kan^R$ | This study |
| pTrc99a | Ptac containing expression vector used for IPTG induced overexpression in *E. coli*, $Amp^R$ | [10] |
| pEK188 | pTrc99a-based plasmid for *aimB*-FLAG expression, $Amp^R$ | This study |
| pBbS2k | Ptet containing low-copy expression vector used for anhydro-tetracycline induced overexpression in *E. coli*, $Kan^R$ | [11] |
| pEK189 | pBbS2k-based plasmid for *aimB*-FLAG expression, $Kan^R$ | This study |
| pUC19 | Plac containing high-copy expression vector used for IPTG induced overexpression in *E. coli*, $Amp^R$ | [12] |
| pEK199 | pUC19-based plasmid with Plac replaced with Ptet-*aimB*-FLAG, $Amp^R$ | This study |
| pEVOL-pBpF | Plasmid encoding a tRNA synthetase/tRNA pair for *in vivo* incorporation of p-benzoyl-l-phenylalanine in *E. coli* amber codons (Addgene #31190), $Chlor^R$ | [13] |
| pZS2-123 | Cloning plasmid with 3 promoters to drive independent expression of CFP, YFP, and mCherry (Addgene #26598), $Kan^R$ | [14] |
| pEK70 | pZS2-123 derived plasmid with CFP cassette removed and mCherry replaced with *aimB*, $Kan^R$ | This study |
| pMreBXL series | pEK70-derived plasmids with YFP replaced by *mreB* containing the respective amber codon mutation; *mreB* mutants synthesized by Genscript (Table S1), $Kan^R$ | This study |
| pMreBXL-14-FLAG | pMreBXL-14-based plasmid with an AimB C-terminal FLAG tag, $Kan^R$ | This study |

**Table S3. Primers used in this study.**

| Name | Sequence |
|---|---|
| *Cloning primers* | |
| pNPTS-F | CTCTGCAGGATATCTGGATC |
| pNPTS-R | CTAGTGAGTCGTATTACGTAG |
| mreB-up-F | cgtaatacgactcactagTGTTCAAGGAACGCCTGACCCCTTTGCAGGTGGTC |
| mreB-up-R | gagccagaGCCGTCGGCCGGCGCGCG |
| msfGFP-F | cgacggcTCTGGCTCGAGCAGTAAAGGTGAAGAAC |
| msfGFP-R | gaccttcGCCCGGCGCGCCAGATTT |
| mreB-down-F | cgccgggcGAAGGTCTGTCGATCGACG |
| mreB-down-R | ccagatatcctgcagagAAGCTTGGGATTGGGCCC |
| aimB-F | tactcatATGACCACCTTCGACGAACG |
| aimB-R | tactgaattcTTACTCAGACTTGATCTGCTCGCG |
| aimB-FLAG-F | gacgacgacaagTAAGCGCTCGGAGCGTCG |
| aimB-FLAG-R | gtccttgtagtcCTCAGACTTGATCTGCTCGCG |
| EK644 | TAGAACTTCCGAGAAGTTCA |
| EK645 | GGTGGTTTGTTTGCCGGATC |
| EK646 | TAATAACCAGGCATCAAATAAAACGAAAGGC |
| EK647 | GGCAGGTGCTCCTTCTTAAAGTT |
| EK648 | tatttgatgcctggttattattaCTCAGACTTGATCTGCTCGC |
| EK649 | agaaggagcacctgccatgACCACCTTCGACGAACG |
| EK679 | TAATAATAACCAGGCATCAAATAAAACGAAAGG |
| EK680 | cttgtcatcgtcatccttgtagtcCTCAGACTTGATCTGCTCGCG |
| EK807 | ATGTATATCTCCTTCTTAAAAGATCTTTTGAATTCTTTTC |
| EK808 | GGATCCAAACTCGAGTAAG |
| EK809 | tttaagaaggagatatacatATGACCACCTTCGACGAAC |
| EK810 | actcgagtttggatccTTACTTGTCGTCGTCGTC |
| EK811 | GGTCTGTTTCCTGTGTGAAATTG |
| EK812 | GGATCCTCTAGAGTCGACC |
| EK813 | acacaggaaacagaccATGACCACCTTCGACGAAC |
| EK814 | cgactctagaggatccTTACTTGTCGTCGTCGTC |
| EK838 | CGAGCTCGAATTCACTGGCC |
| EK839 | CCTGCAGGCATGCAAGCTTG |
| EK840 | agtgaattcgagctcgTTAAGACCCACTTTCACATTTAAG |
| EK841 | ttgcatgcctgcaggTATAAACGCAGAAAGGCC |
| EK1003 | tagtgGCGTTCGTCGAAGGTGGTCA |
| EK1004 | aaacTGACCACCTTCGACGAACGCc |

| QPCR Primers | Forward | Reverse |
|---|---|---|
| *rpoD* | CTCTATGCGATCAACAAGCG | ATAGGCCTTGAGGAACTCGC |
| *aimB* | ACGTGCTGCGCAAGGTCT | CCAGCAGCTCGGCCATTT |

**Table S4. MD simulation systems in this study.**

| Name | Structure source | Ligand | Atoms | Simulation time (ns) | Replicates |
|---|---|---|---|---|---|
| *Cc*MreB | PDB ID: 4CZM | ATP, $Mg^{2+}$ | 84,000 | 100 | 1 |
| *Ec*MreB | PDB ID: 4CZM (homology model) | ATP, $Mg^{2+}$ | 84,000 | 100 | 1 |
| *Cc*MreB - AimB | PDB ID: 4CZM; PDB ID: 2KZC (homology model) | ATP, $Mg^{2+}$, AimB | 89,000 | 100 | 2 |
| *Ec*MreB - AimB | PDB ID: 4CZM (homology model); PDB ID: 2KZC (homology model) | ATP, $Mg^{2+}$, AimB | 89,000 | 100 | 2 |

**Supplemental Figure Legends**

**Supplemental Figure 1: MreB-GFP$^{SW}$ minimally perturbs *C. crescentus* growth and shape.**

A) The growth of wild-type and MreB-GFP$^{SW}$ cells was measured by optical density. Error bars are the standard error (n=3).

B-C) Wild-type and MreB-GFP$^{SW}$ cells were imaged by phase microscopy for quantification of cell width (B) and length (C) (n>70 cells per strain; *p*-values: two-tailed t-test).

**Supplemental Figure 2: Relative expression levels of MreB and AimB.**

A-B) AimB-FLAG and MreB-GFP$^{SW}$ expression were assayed by immunoblotting. Samples were normalized by OD. AimB-FLAG expression was induced in *Caulobacter* with 0.3% xylose for 9 h (A), and in *E. coli* with 1 mM IPTG (Plac) or 100 ng/mL aTc (Ptet) for 9 h (B). Cultures were back-diluted 1:100 after 4.5 h to keep cells in log-phase.

**Supplemental Figure 3: Identification of AimB-overexpression suppressor mutants.**

(A-C) The locations of the identified AimB-overexpression suppressor mutants mapped to structures for AimB (A), MreB (B), and the MreB longitudinal polymerization interface (C). (D) Based on Molecular Dynamics simulations (Figure 4F), MreB residues that come within 5 Å of AimB are highlighted in orange. The protein sequences of *Cc*MreB and *Ec*MreB were aligned using Clustal Omega[15] and the interacting residues are highlighted.

**Supplemental Movies S1-S2: Molecular dynamics simulations of MreB-AimB interactions.**

The interactions between AimB and *CcMreB* (Movie S1) or *EcMreB* (Movie S2) were simulated over 100 ns using Molecular Dynamics (see Experimental Procedures). AimB remains stably associated with *CcMreB* but not *EcMreB*.

Figure S1

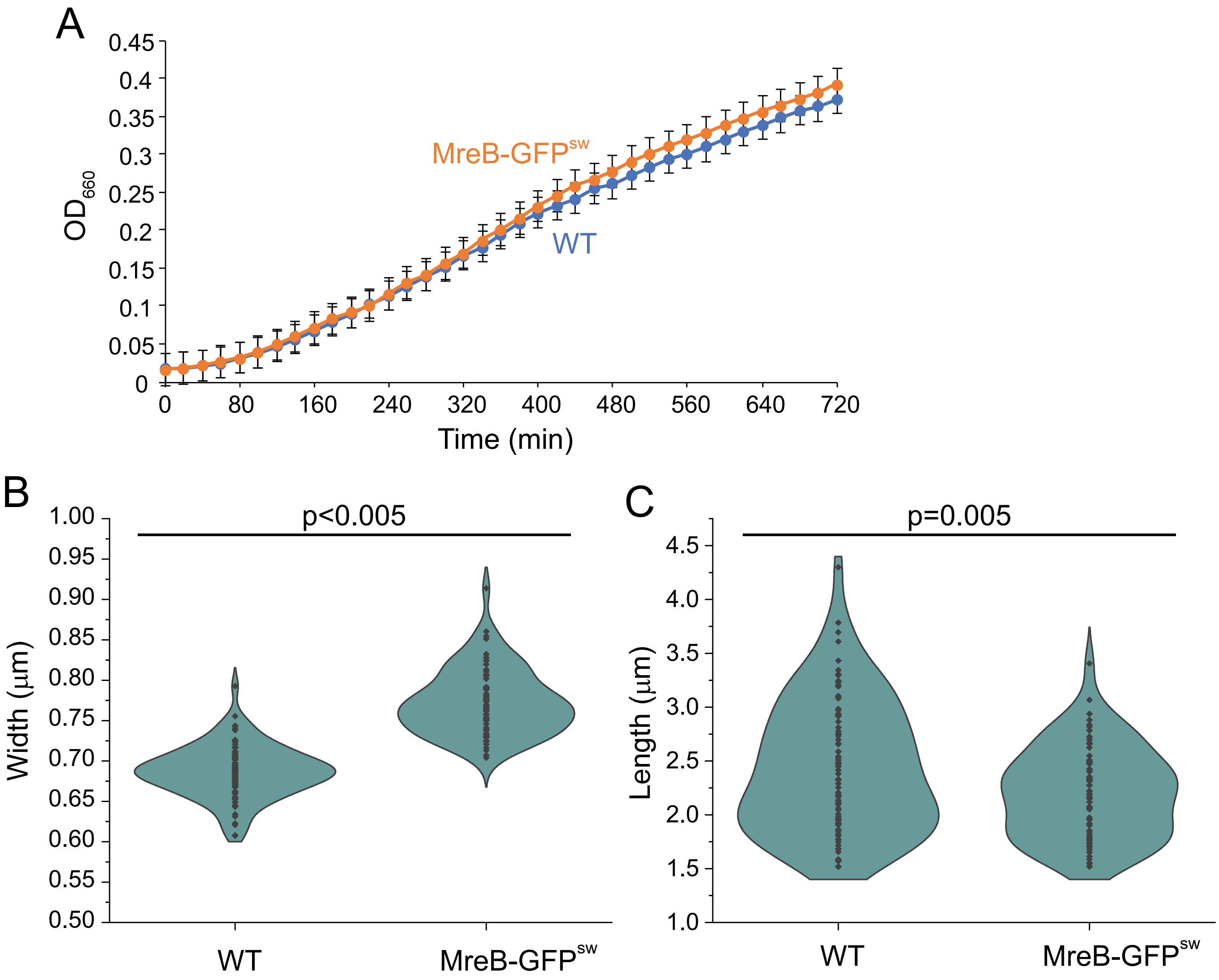

A
0.45
0.4
0.35
0.3
0.25
0.2
0.15
0.1
0.05
0
OD660
MreB-GFPsw
WT
0
80
160
240
320
400
480
560
640
720
Time (min)
B
p<0.005
1.00
0.95
0.90
0.85
0.80
0.75
0.70
0.65
0.60
0.55
0.50
Width (µm)
WT
MreB-GFPsw
C
p=0.005
4.5
4.0
3.5
3.0
2.5
2.0
1.5
1.0
Length (µm)
WT
MreB-GFPsw

Figure S2

A

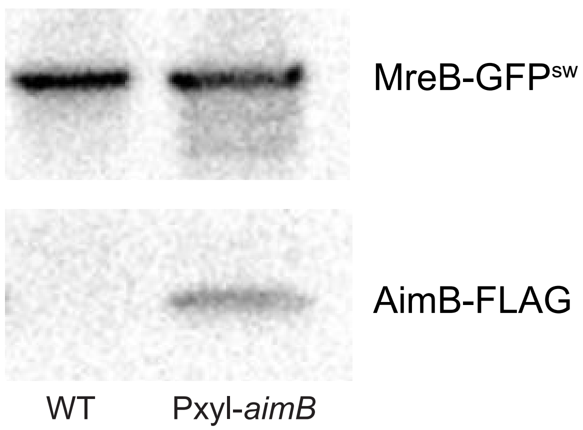
MreB-GFPsw
AimB-FLAG
WT
Pxyl-aimB

B

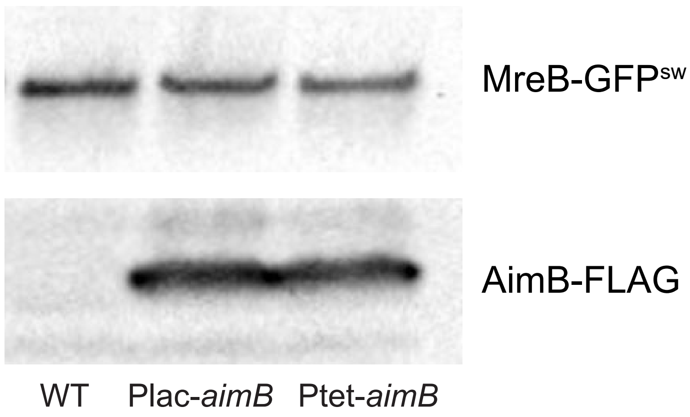
MreB-GFPsw
AimB-FLAG
WT
Plac-aimB
Ptet-aimB

### Figure S3

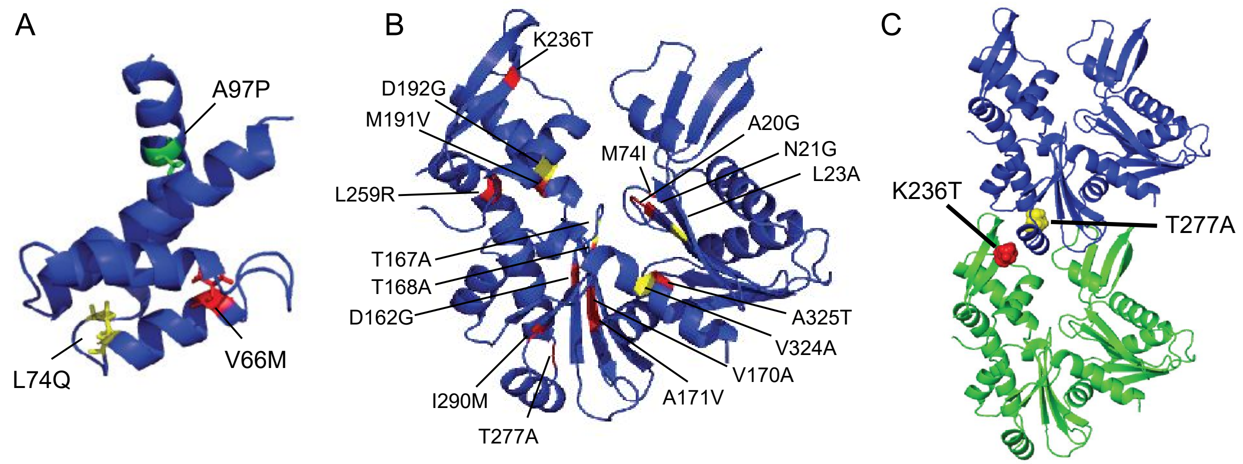


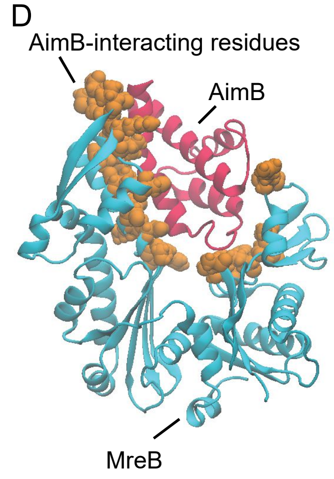


```
CcMreB     MFSSLFGVISNDIAIDLGTANTLIYQKGKGIVLNEPSVVALRN--VGGRKVVHAVGIEAK     58
EcMreB     MLKKFRGMFSNDLSIDLGTANTLIYVKGQGIVLNEPSVVAIRQDRAGSPKSVAAVGHDAK     60
           *:..: *::***::******* **:**********:*:   .*. * * *** :**

CcMreB     QMLGRTPGHMEAIRPMRDGVIADFEVAEEMIKYFIRKVHNRKG-FVNPKVIVCVPSGATA     117
EcMreB     QMLGRTPGNIAAIRPMKDGVIADFFVTEKMLQHFIKQVHSNSFMRPSPRVLVCVPVGATQ     120
           *********:: ******:******* *:*:*:::**::**...    .*:*:**** ***

CcMreB     VERRAINDSCLNAGARRVGLIDEPMAAAIGAGLPIHEPTGSMVVDIGGGTTEVAVLSLSG     177
EcMreB     VERRAIRESAQGAGAREVFLIEEPMAAAIGAGLPVSEATGSMVVDIGGGTTEVAVISLNG     180
           ******.:*. .****.* **:************: * ******************:**.*

CcMreB     IVYSRSVRVGGDKMDEAIISYMRRHHNLLIGETTAERIKKEIGTARAPADGEGLSIDVKG     237
EcMreB     VVYSSSVRIGGDRFDEAIINYVRRNYGSLIGEATAERIKHEIGSAYPG--DEVREIEVRG     238
           :*** *****:***:******.*:**::. *****:******:***:*      .*  .*:*:*

CcMreB     RDLMQGVPREVRISEKQAADALAEPVGQIVEAVKVALEATPPELASDIADKGIMLTGGGA     297
EcMreB     RNLAEGVPRGFTLNSNEILEALQEPLTGIVSAVMVALEQCPPELASDISERGMVLTGGGA     298
           *:* :**** . :..::  :** **:  **.** ****  ********:::*::******

CcMreB     LLRGLDAEIRDHTGLPVTVADDPLSCVALGCGKVLEHPKWMKGVLESTLA     347
EcMreB     LLRNLDRLLMEETGIPVVVAEDPLTCVARGGGKALEMIDMHGGDLFSEE-     347
           ***.**  : :.**:**.**:***:*** * **.**  .   * * *
```
